# Predicting quantitative traits from genome and phenome with near perfect accuracy

**DOI:** 10.1101/029868

**Authors:** Kaspar Märtens, Johan Hallin, Jonas Warringer, Gianni Liti, Leopold Parts

**Affiliations:** Institute of Computer Science, University of Tartu, Tartu, Estonia; Institute of Research on Cancer and Aging, University of Sophia Antipolis, Nice, France; Department of Chemistry and Molecular Biology, Gothenburg University, Gothenburg, Sweden; Centre for Integrative Genetics (CIGENE), Department of Animal and Aquacultural Sciences, Norwegian University of Life Sciences, As, Norway; Wellcome Trust Sanger Institute, Wellcome Trust Genome Campus, Hinxton, United Kingdom

## Abstract

In spite of decades of linkage and association studies and its potential impact on human health^1^, reliable prediction of an individual's risk for heritable disease remains difficult^2-4^. Large numbers of mapped loci do not explain substantial fractions of the heritable variation, leaving an open question of whether accurate complex trait predictions can be achieved in practice^5,6^. Here, we use a full genome sequenced population of 7396 yeast strains of varying relatedness, and predict growth traits from family information, effects of segregating genetic variants, and growth measurements in other environments with an average coefficient of determination *R*^2^ of 0.91. This accuracy exceeds narrow-sense heritability, approaches limits imposed by measurement repeatability, and is higher than achieved with a single replicate assay in the lab. We find that both relatedness and variant-based predictions are greatly aided by availability of closer relatives, while information from a large number of more distant relatives does not improve predictive performance when close relatives can be used. Our results prove that very accurate prediction of heritable traits is possible, and recommend prioritizing collection of deeper family-based data over large reference cohorts.

---

Disease incidence can be predicted based on the health record^7^, the family history^3^, or the genetic risk due to predisposing genetic variants segregating in the population^8^. Each of these sources of information carries signal about the trait, but is not sufficient for accurate prediction. For example, the genetic variants mapped to a trait in genome-wide association studies do not estimate disease risk well, with the vast majority of the heritable variation not accounted for^5,6^, and prediction accuracies lagging far behind narrow sense heritability estimates^9^. An important question of whether this is due to paucity of data, or perhaps more fundamental limitations, can be attacked by predicting phenotypes in model organisms^10,11^. In particular, crosses of founders in the yeast system have circumvented many of the technical difficulties associated with human genetic analyses, and illuminated genetic basis of variation in molecular traits^12-14^, cellular phenotypes^15-17^, missing heritability^18^, and role of interactions^19-21^. For yeast, growth in various environments is an analog of the health record, family history is approximated by phenotypes of closely related individuals, and risk variants can be mapped as for humans. Thus, we can test whether accurate phenotype prediction is possible in practice and what the constraints are.

We used a two-stage crossing design (Fig. S1) to generate 7396 diploid hybrid *Saccharomyces cerevisiae* strains with phased whole genome sequences. Due to the crossing scheme, each hybrid has 170 “close” relatives that share half of its chromosomes, and 7225 “distant” ones, for which no complete chromosome is shared, but a substantial part of linkage blocks and allele combinations are (Fig. S2). We measured their growth in nine environments in technical and biological duplicate, normalized it against hundreds of internal standards, and used the replicate average for analysis. The environments challenge different cellular functions, covering energy sources (e.g. galactose), osmotic stress (e.g. NaCl), and cancer drugs (e.g. rapamycin, Table S1). The phenotype means have large narrow-sense heritabilities (*h*^2^) and repeatabilities (*H*^2^, broad sense heritability) (median *h*^2^ = 80%, *H*^2^ = 94%, standard error = 0.09, Table S2, S3). The traits are not independent (pairwise Pearson’s *r^2^* = 0.01 to 0.49, Fig. S3), reflecting shared genetic, epigenetic, and environmental influences (Fig. S4).

We first test how well different genomic and phenomic data predict growth phenotypes in our population (Fig. 1a, S5), and then combine them using linear mixed models^22^. We obtained predictions via four-fold cross-validation, with the training set randomly sampled from both close and distant relatives (Methods, Fig. S1). One growth trait could be predicted from the rest with reasonable accuracy (Fig. 1b “P”, median *R*^2^ = 0.48), and the quality of prediction depends on the strength of pairwise correlations of the phenotypes. The genomic best linear unbiased predictor (BLUP), an additive model based on realized genetic relatedness alone, captures the pedigree structure in the population, and achieves prediction accuracies very close to the narrow-sense heritability estimates (Fig. 1b “BLUP”, median *R*^2^ = 0.77, 98% of *h*^2^ explained). Interestingly, these predictions are near-identical to a simple midparent approach (Pearson’s *r*^2^ > 0.99, Fig. S6). Thus, the genetic similarity between individuals explains nearly all additively heritable variation in our population.

**Figure 1.**
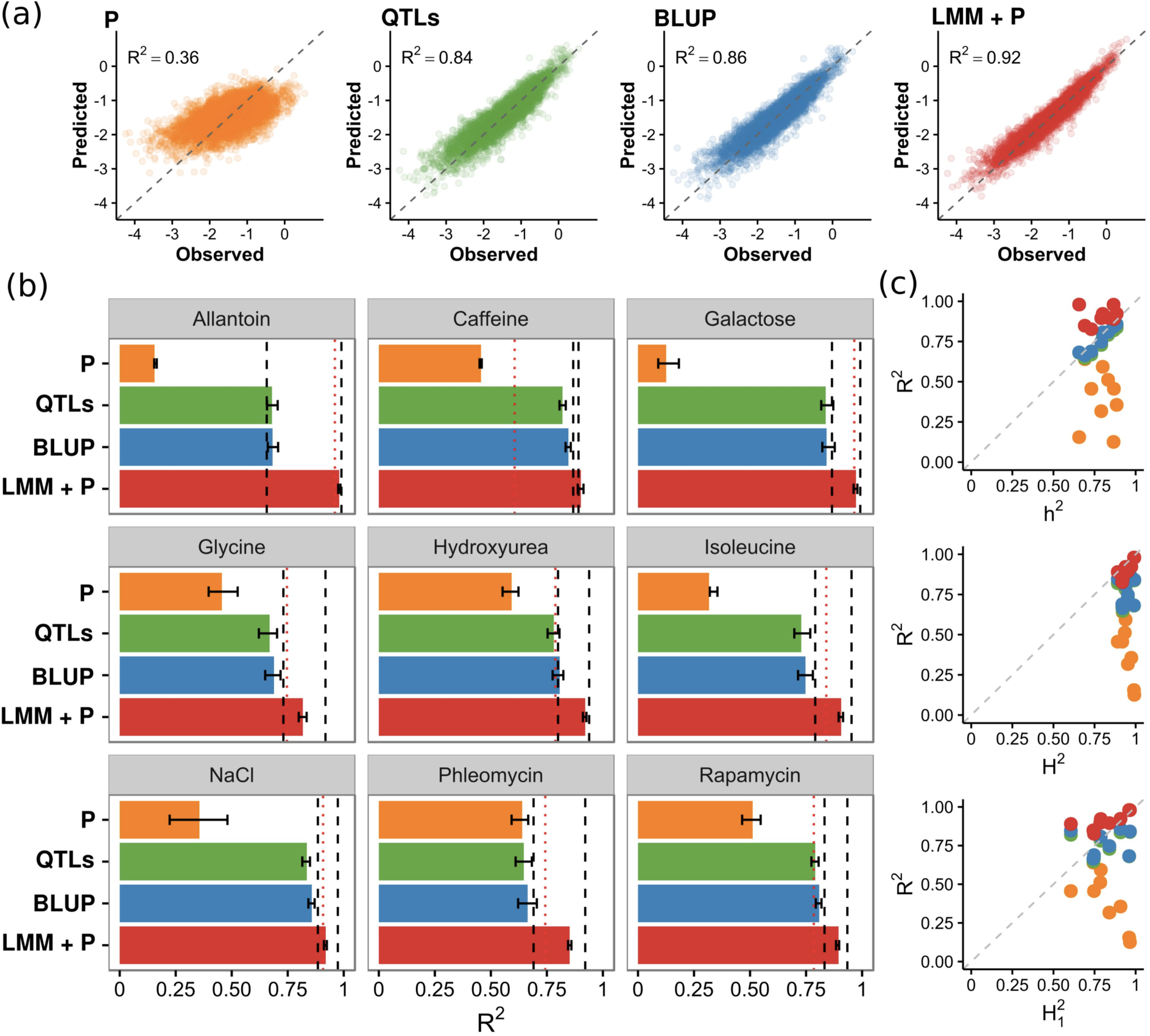
Prediction accuracy. All panels contain four model classes: linear regression on other phenotypes (“P”, yellow), linear regression with additive effects determined by forward selection (“QTLs”, green), prediction based on the realized genetic relatedness (“BLUP”, blue), and the best LMM with additive and interaction effects together with other phenotypes (“LMM+P”, red). All prediction accuracies denote coefficient of determination R, and are determined by four-fold cross-validation. (a) Models using a single source of information predict less accurately than a combined one. Predicted (*y*-axis) and observed (*x*-axis) growth in NaCl for every measured hybrid strain (dots) for each model class, with coefficient of determination (*R*^2^) of the predictions labeled. Perfect predictions would lie on the grey dashed line *y*=*x*. (b) Linear mixed models with information from other phenotypes give very accurate predictions. Predictive performance (*R*^2^, *x*-axis) for different models (*y*-axis) for each of the measured phenotypes (nine boxes). Bars indicate therange of *R*^2^ over the four cross-validation folds. The dashed lines show narrow-sense heritability *h*^2^ (black, left) and repeatability *H*^2^ (black, right) estimates for the mean phenotype, and the dotted line (red) shows repeatability of a single measurement 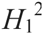. (c) Prediction can be more accurate than one measurement. Prediction accuracy of mean phenotype (*R*^2^, y-axis) compared to different types of heritability estimates (*x*-axis) for the four model classes: narrow-sense heritability of average phenotype (*h*^2^, top panel), repeatability of average phenotype (*H*^2^, middle panel), and repeatability of a single measurement (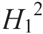, bottom panel). Grey dashed lines denote the identity *y*=*x*.

Next, we mapped quantitative trait loci (QTLs) in each environment, and asked how well they predict growth in that environment. A small number of SNPs with the largest effects explain a sizeable portion of additive variance, but for all traits the prediction accuracy remains lower than BLUP’s (e.g. median *R*^2^ = 0.58 vs 0.81 for 10 QTLs, Fig. S7). When up to 50 SNPs are included in the model, the accuracy reaches *h*^2^ (Fig. 1b, “QTLs”, median *R*^2^ = 0.78, 98% of *h*^2^ explained), with predictions very similar to BLUP (*r*^2^ > 0.97, Fig. S8). Therefore, all tested methods that consider additive genetic effects reach the same, near-*h*^2^ performance, and there is no missing narrow sense heritability in our experiment. Extending to the linear mixed model (LMM) framework to include genetic background, dominance and interaction effects gave a modest further improvement (median increase of *R*^2^ by 0.03), mainly due to dominance effects of strongest QTLs for allantoin and galactose (Fig. S9).

We then included other phenotypes measured for the same individual as covariates in the model, and achieved median prediction accuracy of 0.91 (Fig. 1b “LMM+P”). To our knowledge, this is the highest for complex traits to date^23,24^, exceeding narrow-sense heritability for all nine phenotypes, and approaching repeatability (Fig. 1c, 96% of *H*^2^ explained). For each of the measured traits, our predictions of the mean phenotype (i.e. the average of four replicate measurements) have lower error than a single growth experiment (Fig. 1c). The combined model improves over others especially when a large proportion of heritable non-additive variation is not captured by interaction and dominance effects (Fig. S4). We also applied the recently published mixed random forest approach, which accounts for population structure and captures non-linear genetic effects^25^. This method performed similarly to the combined LMM+P model (median *R*^2^ 0.91 vs. 0.91), with no consistent difference across the traits (Fig. S10).

So far, our predictions for each test individual were obtained from models that were trained with data from its close relatives. We observed that errors were larger when close relatives were not available (e.g. Fig. 2a, Fig. S11). Thus, we next compared two training scenarios - “close relatives”, where each member of the test set has several close relatives in the training set, and “distant relatives”, where test set individuals are not closely related to anyone in the training set (Fig. S1b). When training on close relatives, predictions based on other traits of the same individual are slightly more accurate (median improvement = 0.04, Fig. 2b, “P”), whereas BLUP performs substantially better. On average, BLUP achieves *R*^2^ of 0.14 when trained on distant relatives and 0.76 on close ones (Fig. 2b, “BLUP”). This difference is explained by the larger uncertainty of the predictive distribution based on distant relatives: the observed errors are near-perfectly calibrated to their model-derived standard errors (Fig. 2c, *r*^2^=0.96). Accuracy increases markedly even with a small number of close relatives included in the training data, while adding more distant relatives to close ones does not improve predictions (Fig. 2d, Fig. S12). For example, adding on average just five close relatives per test individual rises the median *R*^2^ from 0. 15 to 0.65, but complementing the training set of close relatives by all distant relatives has a negligible effect (median *R*^2^ = 0.79 vs. 0.81).

**Figure 2.**
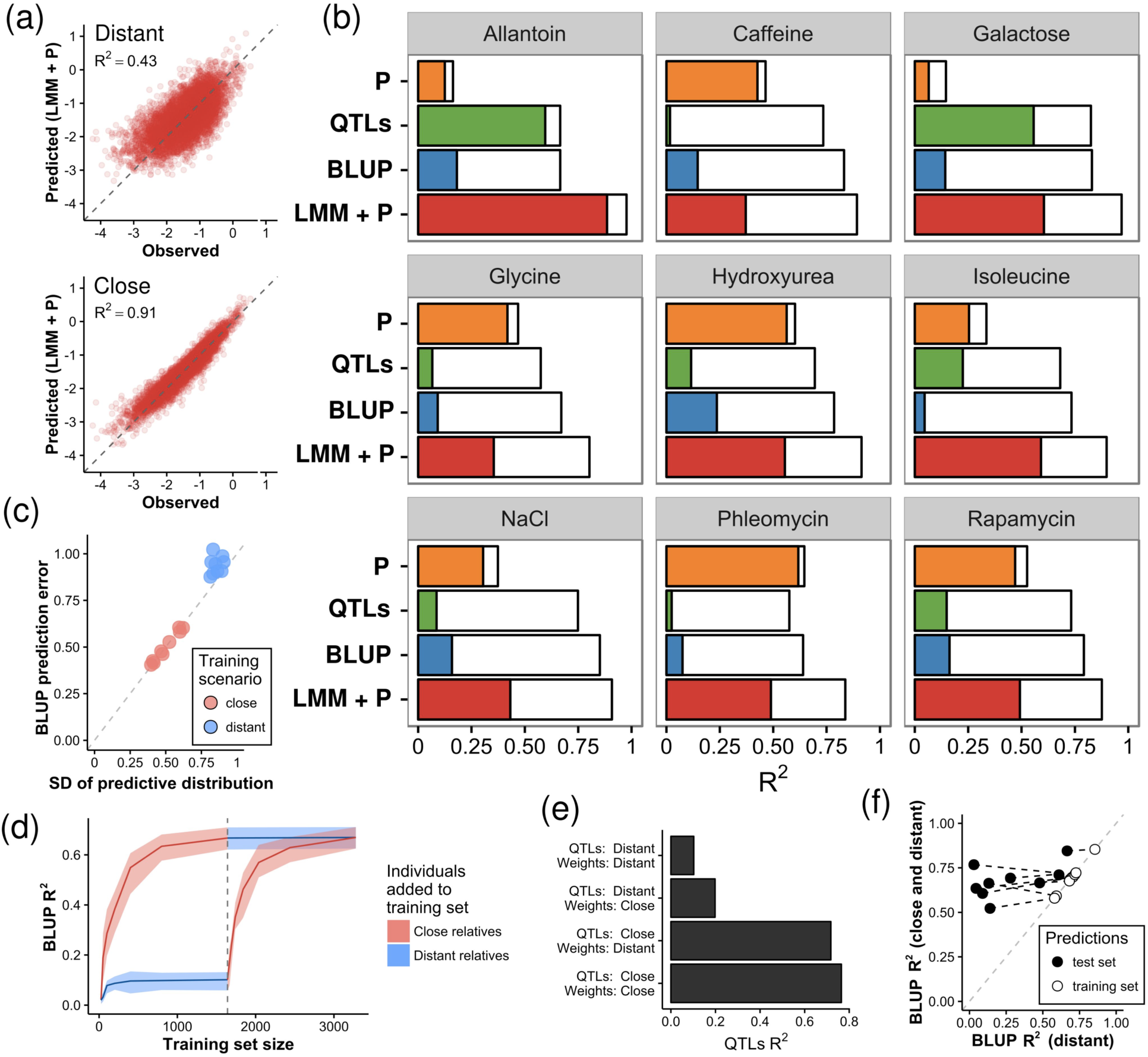
Close relatives improve predictions. All panels cover two scenarios: training on “close relatives”, where each member of the test set has several close relatives in the training set, and “distant relatives”, where test set individuals are not closely related to anyone in the training set. Predictions are obtained by four-fold cross-validation. (a) Close relatives greatly contribute to genome-based prediction accuracy. Predicted (*y*-axis) and observed (*x*-axis) growth for test set individuals (blue dots) in NaCl using the best LMM+P model in different training scenarios. Grey dashed line denotes the identity *y=x*. (b) Distant relatives are more difficult to predict in each environment. Predictive performance (*R*^2^, *x*-axis) of different model classes (*y*-axis) in two training scenarios: “Distant” (colored bars) and “Close” (white bars). (c) BLUP predictions from distant relatives are less accurate due to a more uncertain model-derived predictive distribution. Prediction error (*y*-axis, standard deviation of the residuals) compared to the standard deviation of the predictive distribution (*x*-axis) for the nine environments, when trained on distant (blue dots) or close relatives (red dots). (d) BLUP predictions are more accurate when the model is trained on a small number of close relatives compared to a large set of distant relatives. Predictive performance of BLUP (*R*^2^, *y*-axis) improves with expanding the training set (size on x-axis) with individuals closely (red line) or distantly (blue line) related to the test set. From the dashed grey line onwards, distant relatives are added to the training set of closely related individuals, and vice versa. Shaded regions denote the range of *R*^2^ over the four cross-validation folds. (e) Low QTL predictive ability for out-of-sample distant relatives is mainly due to discrepancies between the sets of mapped QTLs, not their estimated effects. Predictive performance (*R*^2^, *x*-axis) of the QTLs model, stratified by training sets used for QTL mapping (model selection) and weight estimation (model fitting). QTL mapping and weight estimation are carried out under four training scenarios (*y*-axis): both stages in distant relatives (“QTLs: Distant, Weights: Distant”), both in close relatives (“QTLs: Close, Weights: Close”), QTLs mapped in distant relatives and weights estimated in close relatives (“QTLs: Distant, Weights: Close”), or vice versa (“QTLs: Distant, Weights: Close”). (f) A minor change in the training set (replacing 1% of distant relatives with close ones) has a profound effect on out-of-sample QTL-based prediction accuracy. Out-of-sample (black dots) and in-sample (white dots) predictive performance (*R*^2^) of QTLs model in two scenarios: trained on distant relatives only (*x*-axis) or when 1% is replaced with close relatives (y-axis).

Perhaps surprisingly, training on close relatives improved also QTL-based predictions. For near-monogenic traits (e.g. growth in allantoin and galactose), the accuracies were similar for both training scenarios (Fig. 2b “QTLs”). However, for more complex traits, the QTL model trained on distant relatives reaches high accuracy in the training data, but does not perform well out of sample, with 61% median decrease in accuracy (respective decrease for close relatives is 3%, Fig. 2c). In this case, the prediction uncertainties are similar (Fig. S13), and most of this difference is explained by model selection. When we mapped QTLs in close relatives, but estimated their weights on distant relatives, the prediction accuracy decreased from 0.73 to 0.65 compared to carrying out both procedures on close relatives (Fig. 2e, Fig. S14). Conversely, mapping QTLs in distant relatives and fitting their weights in close relatives resulted in a much lower *R*^2^ of 0.31. Including close relatives in training gives a more robust approximation of the phenotypic covariance structure (Fig. S15), which explains the large gap between out-of-sample and in-sample performance for distant relatives (Fig. 2f). Notably, prediction accuracy drops substantially, even when just 1% of the training data changes (Fig. 2f).

Combining genomic and phenotypic information (LMM+P) to predict from distant relatives gives accuracies similar to combining QTLs and phenotypic information. For traits where genomic prediction on distant relatives does not work well (e.g. caffeine, phleomycin), this model performs similarly to using other phenotypes only or even slightly worse (median improvement 0.02, Fig. 2b “LMM+P”). However, for traits with large effect QTLs (allantoin, galactose, isoleucine), genetic information helps prediction even if BLUP is not accurate.

We predicted nine heritable traits in a population of 7396 yeast strains of varying relatedness, and achieved accuracies over 90%, very near the repeatability limit. To our knowledge, these are the most accurate out-of-sample predictions of complex traits to date. There is almost no missing narrow or broad sense heritability, proving that very accurate genome-aided predictions can be obtained in practice, in contrast to relatively poor genomic prediction performance for human cohorts, e.g. *R*^2^ < 0.16 using distant, and < 0.37 for close relatives^9^. The improvement in predictive ability beyond *h*^2^ using phenotype data is due to capturing additional signal from the environmental and non-additive genetic components, reflecting the extent to which these are shared between the traits.

When no close relatives were available, and no single QTL explained a large fraction of variance, the pure genomic methods were inaccurate, even in our population of over 7000 individuals. At the same time, when the number of close relatives in the training sample was sufficiently large, the predictions were not improved by adding all remaining distant relatives. In concert, these observations suggest that efforts directed towards creating genotype-based scores using common variants to predict disease risk would benefit dramatically from being complemented by systematic collection of family history and relatedness data^26,27^. Since information from as few as five close relatives gave large gains, we expect such an approach to be a cost-effective solution for achieving better prediction in a clinical setting with finite resources.

## Acknowledgements

KM was supported by the European Regional Development Fund through the BioMedIT project, JH by the LABEX SIGNALIFE (ANR-11-LABX-0028-01), Swedish Research Council (grant numbers 325-2014-6547 and 621-2014-4605) and the Research Council of Norway (grant number 222364/F20), JH and GL by ATIP-Avenir (CNRS/INSERM), ARC (grant number SFI20111203947), FP7-PE0PLE-2012-CIG (grant number 322035), ANR (ANR-13-BSV6-0006-01), and Canceropole (AAP, Labex 11-LABX-0028-01), and LP by a Marie Curie International Outgoing Fellowship, the Wellcome Trust, and Estonian Research Council (IUT34-4). We thank Francisco Salinas for technical help with large-scale crossing, Martin Zackrisson for much appreciated technical assistance with extraction and analysis of growth estimates, and Oliver Stegle, Cornelis Albers, and Daniel Gaffney for comments on the text.

## Methods

### Panel design, genotyping, phenotyping, normalization

172 haploid F_12_ segregants (86 MatA and 86 MatAlpha) from a cross between YPS128 and DVPBG6044^28^ were crossed in an all against all fashion to obtain 86 × 86 = 7396 diploid hybrids using standard yeast protocols (Fig. S1). The strains were grown in biological and technical duplicates (four measurements total) in 1536-position solid agar plate cultures, where every fourth position was the reference YPS128 strain for normalization purposes. We extracted endpoint growth (i.e. final number of cells) in each of the nine environments, converted it to logscale, and normalized to the internal YPS128 control. The four replicate values were then averaged to obtain the final phenotype (i.e. mean growth) for each individual and environment.

### Modelling and predictions

We used a range of models to predict a trait of interest either on genomic information only, individual phenotypic information only or both.

**Phenotype (“P”)**. Let *y* be the vector containing the phenotype of interest for all *N* individuals, and let *P*_1_*,…, P*_8_ be the remaining phenotypes. We modelled y as *y* ~ *N*(*β*_0_ *+β*_1_*P*_1_+ … +*β*_8_*P*_8_, *σ*^2^*I*) to fit the phenotype weights β used for prediction.

**Best linear unbiased predictor (“BLUP”).** Let *x_j_* be the genotype vector for SNP *j=1, …, M,* and let *X* be the genotype matrix *X=(x_1_, …, x_M_).* In the genomic BLUP model, *y* = *μ*1 + Σ*_j_ b_j_x_j_ + ε* with random coefficients 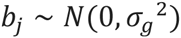 and measurement noise 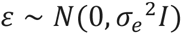. This model implies the multivariate Gaussian distribution 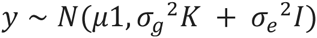, where 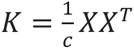 is the realized genetic relatedness matrix, with the scaling constant *c* being the average diagonal value of *XX^T^*. Prediction for the test individual can be obtained by conditioning on the observed data in a standard way for multivariate normal distributions. When calculating the standard deviation of the predictive distribution (Fig. 2c), we averaged the variances on the predictive distributions (i.e. averaged the diagonal elements of the covariance matrix of the predictive multivariate normal distribution) and reported the square root of this number.

**Quantitative trait loci (“QTL”)**. To identify the strongest QTLs, we first carried out forward selection for up to 50 iterations in the linear regression model *y* ~ *N*(*β*_0_ *+Σ,j_ϵQt_ β_j_x_j_, σ*^2^*I*), where *Q_t_* denotes the selected collection of QTL indexes at iteration t. The number of QTLs in the final model was determined by out-of-sample prediction accuracy, with four-fold cross-validation on the training portion of data (hence, altogether a double cross-validation scheme).

**Midparent**. Let *y_ij_* be the phenotype for individual who has parents *i* and *j*. Let 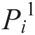 and 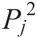 be the parental phenotype values. We model *y_ij_* as the midparent value 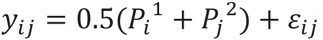 where *ε_ij_* is uncorrelated noise. We first fit the parental values from the *y_ij_* observed in training data, and used them to predict phenotypes of test individuals.

**Linear mixed model including phenotypes (“LMM+P”).** The LMM+P model combines additive, dominance and interaction effects with genetic relatedness and other traits, 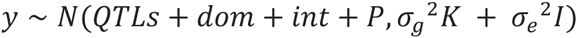. The fixed effects contains of a genetic (*QTLs+dom+int*) and non-genetic (*P*) part. The latter includes the linear combination of all other traits *P_1_, …, P_8_.* The genetic component is constructed with forward selection among additive QTLs and interaction between all such SNP pairs *x_i_* and *x_j_*, where *x_i_* has previously been selected into the model. While we miss interactions where neither locus has a significant additive effect, it has been shown that such occurrences are rare^29^, and their contribution to explaining variance is negligible^19^. By allowing self-interactions, we also incorporated dominance effects. We selected the final model by performing cross-validation on training data after each of the feature selection steps.

### Training and obtaining predictions

All models were fitted with the Python package LIMIX^22^. We used four-fold cross-validation to obtain out-of-sample predictions for all *4N*^2^=7396 individuals by splitting the *4N*=172 parents into four groups of size *N* each, resulting in four subsets of size *N*^2^, and in turn, using one of these as a test set to obtain predictions and the remaining three as a training set to fit the models. First, we did not take into account the relatedness structure and divided individuals into subsets randomly (results in Fig. 1). Later, we distinguished between closely and distantly related individuals (results in Fig. 2). The four test sets remained the same as before, but instead of training on all 3*N*^2^ remaining individuals, we picked the *N* × *N* individuals who do not share a parent with anyone in the test set (“distant relatives”), as well as sampled *N*^2^ from the 2*N*^2^ remaining individuals who do share one parent with someone in the test set (“close relatives”).

### Heritability estimation

Narrow-sense heritability was estimated from the genomic BLUP model as 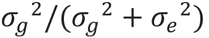, when fitted to all of the data. To estimate repeatability, we fitted the following fixed effects model *r_ij_ = y_i_ + ε_ij_* where *r_i1_, r_i2_, r_i3_, r_i4_* are the four replicate measurements for individual *i*, *y_i_* is the average *r_ij_* value for this individual, and *ε_ij_* ~ *N*(0, *σ*^2^). Repeatability was estimated as 1 − σ^2^/Var(*r*).

